# Mixed Waste Contamination Selects for a Mobile Genetic Element Population Enriched in Multiple Heavy Metal Resistance Genes

**DOI:** 10.1101/2023.11.17.566018

**Authors:** Jennifer L. Goff, Lauren M. Lui, Torben N. Nielsen, Farris L. Poole, Heidi J. Smith, Kathleen F. Walker, Terry C. Hazen, Matthew W. Fields, Adam P. Arkin, Michael W. W. Adams

**Author notes:** **Corresponding Author:** Michael W. W. Adams, Department of Biochemistry & Molecular Biology University of Georgia, Athens, GA 30602, USA.

## Abstract

Mobile genetic elements (MGEs) like plasmids, viruses, and transposable elements can provide fitness benefits to their hosts for survival in the presence of environmental stressors. Heavy metal resistance genes (HMRGs) are frequently observed on MGEs, suggesting that MGEs may be an important driver of adaptive evolution in environments contaminated with heavy metals. Here, we report the meta-mobilome of the heavy metal contaminated regions of the Oak Ridge Reservation (ORR) subsurface. This meta-mobilome was compared to one derived from samples collected from unimpacted regions of the ORR subsurface. We assembled 1,615 unique circularized DNA elements that we propose to be MGEs. The circular elements from the highly contaminated subsurface were enriched in HMRG clusters relative to those from the nearby unimpacted regions. Additionally, we found that these HMRGs were associated with Gamma and Betaproteobacteria hosts in the contaminated subsurface and potentially facilitate the persistence and dominance of these taxa in this region. Finally, the HMRGs were associated with conjugative elements, suggesting their potential for future lateral transfer. We demonstrate how our understanding of MGE ecology, evolution, and function can be enhanced through the genomic context provided by completed MGE assemblies.

## INTRODUCTION

An environmental mobilome, or meta-mobilome, consists of all mobile genetic elements (MGEs) found within an environmental sample ^1^. MGEs are pieces of DNA that can move within the genome of an organism or between the genomes of two different organisms. Prokaryotic MGEs include transposable elements, plasmids, and viruses/phages. MGEs are the major vehicle for movement of genetic material between prokaryotes, a process known as horizontal gene transfer (HGT) ^2^. The spread of MGEs within microbial communities allows for acquisition of novel traits that may facilitate adaptation to fluctuating resources and environmental stressors, driving the ecological diversification of close microbial relatives ^3^. Common MGE-encoded traits include novel carbon substrate utilization ^4^, antibiotic resistance ^5^, toxin production ^6^, and heavy metal resistance ^7^.

Anthropogenic compounds that induce cellular stress responses can increase rates MGE mobilization in microbial populations ^8^. Thus, heavily polluted environments may be major hot spots of HGT. The Oak Ridge Reservation (ORR), located in Oak Ridge, TN, is a well-characterized experimental site for examining the ecological impacts of legacy industrial waste^9^. The near-source subsurface is highly acidic and contaminated with a mixture of nitrate, uranium, and other heavy metals (*e.g.,* Ni, Co, Cu, Fe, Al, Cd, Mn, Hg). Prior studies of ORR microbial populations have suggested that their adaptive evolution may have been facilitated by the lateral transfer of MGEs. For example, metal efflux pumps and mercury resistance genes were shown to be highly mobilizable among *Rhodanobacter* species that predominate in the contaminated ORR groundwater ^10,11^. Martinez, et al. ^12^ found evidence for horizontal transfer of heavy metal-translocating P-type ATPases among a large collection of isolates from the ORR.

High-throughput sequencing has enabled the untargeted, culture-independent analysis of MGEs from diverse environments^13–17^. However, only a subset of these studies reported circularized MGEs. Recently, using methodology to enrich for plasmids during DNA extractions, Kothari, et al. ^18^ recovered 615 circularized MGEs from an aquifer and Kirstahler, et al. ^19^ recovered 159,322 circularized MGEs from global sewage samples. However, in the latter case, the circularized MGEs skewed towards shorter sizes (all < 17,400 bp). Similarly, Kothari et al. recovered a relatively small number of MGEs >20,000 bp in length (5.7 %, 35/615 circular elements) with only three (0.5%) being >100,000 bp in length. These results may represent a filtering effect of the targeted enrichments for MGEs from environmental samples. We suggest that these prior studies fail to adequately capture the diversity of mid-(∼20,000 – 100,000 bp) to larger-sized (≥ 100,000 bp) MGEs that are known to be widely distributed in microbial populations^20,21^, in part, due to the challenges of plasmid assembly from metagenomics data. However, assembling complete, circular MGEs from metagenomes without enrichment is challenging because (1) MGEs are often in low abundance and (2) MGEs often share sequences with bacterial genomes and other MGEs ^22^. Their low abundance means that the MGEs may not be fully covered by short-read sequencing data. The shared sequences result in complex and tangled assembly graphs, resulting in fragmented assemblies ^23^.

The cataloguing of MGE gene function predictions and host range is essential to accurately model microbial species abundances, population dynamics, and functional output of environmental microbiomes at the ecosystem level l^24^. Thus, there is a continued need for method optimization for recovery of diverse MGEs across a representative range of sizes from metagenomes. Here, we use SCAPP, or Sequence Contents-Aware Plasmid Peeler, an algorithm designed for the specific purpose of reconstructing plasmid sequences from metagenomic data^22^. SCAPP leverages biological knowledge of plasmid sequences and uses plasmid-specific genes to annotate the assembly graphs. Nodes in the graph are assigned weights based on the likelihood that they represent plasmids. SCAPP prioritizes peeling off circular paths in the assembly graph that include plasmid genes and highly probable plasmid sequences. It also utilizes plasmid specific genes and plasmid scores to filter out potential false positives. Despite its initial design for plasmid circularization, SCAPP is also very good at circularizing other MGEs, likely due to the genetic similarities between all MGEs ^25^ and its ability to extract circular DNA elements from assembly graphs. We hypothesized that distinct populations of MGEs would be present in the high and low contamination regions of the ORR subsurface. We further predicted that MGEs from the highly contaminated regions of the ORR subsurface would be enriched in clusters of multiple heavy metal resistance genes due to their role in the adaptive evolution of microorganisms in these anthropogenically-disturbed environments.

## MATERIALS AND METHODS

### Sampling and geochemical data

The majority of the sub-surface well geochemistry data were obtained from publicly available datasets, either from Smith, et al. ^26^, Wilpiszeski, et al. ^27^, Gushgari-Doyle et al. ^28^, or from the publicly accessible U.S. Department of Energy Office of Science Subsurface Biogeochemical Research web page (https://www.esd.ornl.gov/orifrc/)^29^. For this study, we performed additional mercury analyses by ICP-MS (**Supplemental Methods**). Not all well samples collected for metagenome sequencing had a complete set of associated geochemical metadata. However, uranium (U) measurements were available for all samples and were used to distinguish between high and low contamination sites. We divided the sampling sites into two categories based on the U.S. Environmental Protection Agency (EPA) maximum contaminant level (MCL) U in drinking water ^30^: (1) highly contaminated sites with [U] > 0.126 µM and (2) low contamination sites with [U] < 0.126 µM). For the sediment samples, contamination levels were inferred from the [U] of the groundwater of adjacent wells.

### Metagenome sequencing and mobile genetic element assembly

Samples for metagenome sequencing and assembly were collected from well water pumped from 17 sites across the Oak Ridge Reservation located within the Bear Creek Valley in Oak Ridge, TN, USA **(Fig. 1A, Table S1, S5)**. At three of these sites (FW306, EB106, and EB271), sediment cores were also collected. Metagenome sequencing and assembly methods are described in Tian et al 2020, Wu et al 2023, and Lui et al (MRA). Reads for EB106, EB271, GW271, FW115-24 metagenomes were deposited in NCBI’s Sequence Read Archive in BioProject PRJNA1001011 under accession numbers SAMN36786281-SAMN36786357. The assembly graphs and sequencing reads were used as input into SCAPP^22^ using default parameters to obtain circular DNA elements. The circular elements were de-replicated within the high and low [U] sample sets prior to further analyses.

**Figure 1.**
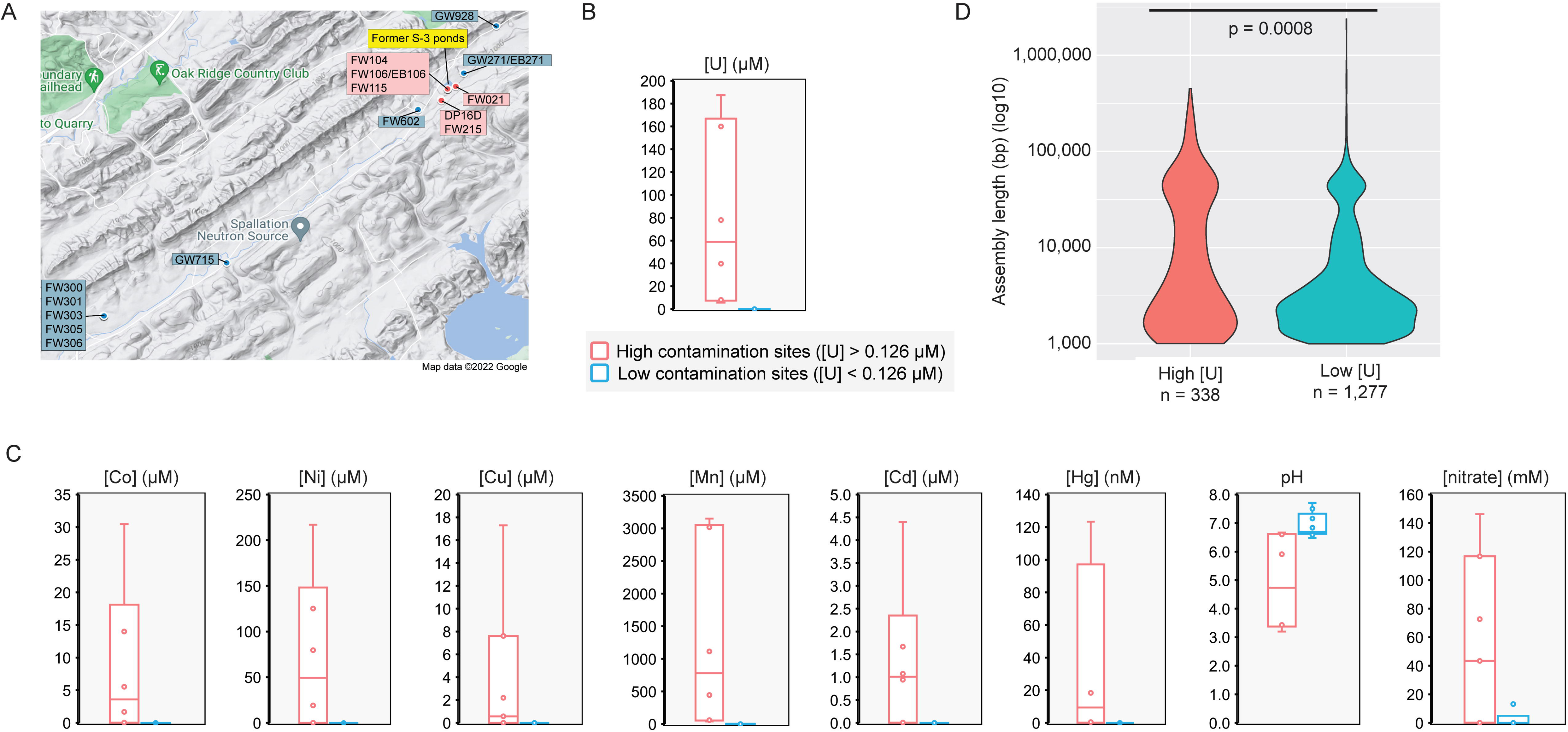
Origin geochemistry and size distribution of ORR MGEs. For all panels red colors indicate a high [U] area while blue colors indicate a low [U] area (A) Map showing the sampling locations within the ORR. (B) Distributions of uranium concentrations between high and low [U] regions of the site. (C) Distributions of other metal concentrations, nitrate concentrations, and pH between high and low [U] regions of the site. (D) Violin plots of the size distribution of dereplicated MGE assemblies between the high and low [U] regions of the site. A Welch’s two-sided t-test was used to test for significant differences between the two distributions.

### Circular element annotations

Annotation of the circular elements was performed using eggNOG-mapper v2 with default parameters ^31^ and the Annotate Domains in a Genome (v1.0.10) application in the Department of Energy (DOE) KnowledgeBase (KBase) with default parameters. This program identifies protein domains using the following domain libraries: Clusters of Orthologous Genes (COGs) ^32^, NCBI’s conserved domains database (CDD models)^33^, Simple Modular Architecture Research Tool (SMART)^34^, NCBI’s Protein Clusters models ^35^, TIGRFAMs hidden Markov models^36^, Pfam hidden Markov models, and NCBI Prokaryotic Genome Annotation Pipeline (PGAP)^37^ hidden Markov models.

### Classification of circular elements

Circular elements representing viral genomes were predicted using VirSorter (v1.0.5) ^38^ implemented in the Department of Energy (DOE) KnowledgeBase (KBase) ^39^. Low confidence category 3 and 6 predictions were removed from the analysis. These circular elements are referred to as “viral genomes” in the text. Taxonomic classification of categories 1, 2, 4, and 5 assemblies was performed using vConTACT2 (v0.9.19) ^40^ implemented in KBase with default parameters. PhaGCN with default parameters was utilized for viral taxonomy classification based off ICTV2022 ^41,42^. PhaTYP with default parameters was utilized for phage lifestyle prediction^43^. We compared the remaining non-viral circular elements to known plasmids in the Plasmid Database (PLSDB, v2021_06_23_v2) using the default search parameters for *mash dist, p = 0.1, distance = 0.1.* The circular elements with similarity to known plasmids are referred to as “plasmids” in the text. The circular elements that were not already classified by PLSDB or VirSorter were further analyzed by searching the eggNOG-mapper annotation files for mobile-genetic element associated Pfams **(Table S4)**. These circular elements are referred to as “Unclassified MGEs” in the text. For the remaining circular elements, we manually examined the annotation files from the comparisons to CDD models, SMART, NCBI’s Protein Clusters models, TIGRFAMs, Pfam hidden Markov models^44^, and PGAP hidden Markov models. This allowed us to identify additional “Unclassified MGEs”. All remaining circular elements are referred to as “cryptic circular elements” in the text.

### Host prediction

Viral hosts were predicted using two parallel methods: *(i) k-mer similarity*: as a broader primary method, hosts for all assemblies were predicted using the Prokaryotic Virus Host Predictor (PHP) Tool^45^. PHP has an accuracy of 80% at the phylum level. These phylum-level assignments were used for further analysis. *(ii) CRISPR spacer matches*: A BLASTn search of viral assemblies was performed against the IMG/VR v4 databases of isolate and uncultivated microorganism CRISPR spacers ^46^. Default parameters were used. The results were filtered to allow for 0 or 1 mismatches between the sequences.

Phylum- and class-level taxonomic assignments were performed for the non-viral circular elements using a gene taxonomy-based approach. Phylogenetic assignments for each annotated coding sequence were extracted from the eggNOG-mapper annotation files. The phylogenetic assignments were tabulated for each non-viral assembly. A taxonomic assignment at the phylum level was made if >50% of CDS belonged to the same phylum or class. A similar analysis was performed for class-level predictions. For all downstream analyses, viral hosts were considered separately from non-viral hosts and different taxonomic levels were analyzed independently.

### Functional analyses

To examine heavy metal resistance and antibiotic resistance genes on the circular elements, Clusters of Orthologous Genes (COGs) annotations were extracted from the eggNOG-mapper annotations. From the COG database, we curated lists of heavy metal resistance and antibiotic resistance-related COGs **(Table S2)**. We then searched the annotation files for these curated COGs. These COG counts were normalized against the total number of coding sequences (CDS) found on the circular elements in each sample set (*i.e.,* high [U], low [U] sets). For identification of conjugative elements, we curated a list of conjugative transfer system COGs and searched the eggNOG-mapper annotation files for these COGs **(Table S3).** Normalization was performed by the same method described above. Functional enrichment calculations were performed using a two-tailed Fisher exact test.

### Data visualization

The following R (v. 4.3.0) packages were used: *circlize* (v0.4.15) ^47^, *pheatmap* (v1.0.12)^48^, and *ggplot2* (v.3.4.2)^49^. Plasmid maps were generated in Geneious Prime (v2022.2). Network diagrams were generated using Cytoscape (v 3.9.1) ^50^.

## RESULTS

### Metagenome-assembled putative mobile genetic elements from the Oak Ridge Reservation subsurface

From a total of 32 sediment and 25 groundwater metagenomes from the Oak Ridge Reservation (ORR)^51–53^, we input the assembly graphs into SCAPP to obtain 1,713 unique, circular elements (>1000 bp in length) representing putative mobile genetic elements (MGEs)^54^. We divided these circular elements into two contamination levels: (1) seven high [U] and (2) ten low [U] sites (**Fig. 1A, B, Table S1)**. For the sediment samples, contamination levels were inferred from the groundwater of adjacent wells. Compared to low [U] sites (median = 0.0 µM), the high [U] (median = 58.9 µM) had higher levels of other metal contaminants including manganese (Mn) cobalt (Co), nickel (Ni), copper (Cu) cadmium (Cd) and mercury (Hg) as well as higher concentrations of nitrate and lower pH **(Fig. 1C)**.

We manually examined the annotation files of these circular elements to identify false positive MGEs, removing 98 assemblies that included (1) partial mitochondrial/chloroplast genomes or (2) likely long repeat regions that were inappropriately circularized. This resulted in 338 putative MGEs from high [U] sites and 1,277 putative MGEs from low [U] sites **(Table S6)**. Out of the 1,615 circular elements, 259 (16%) were greater than 20,000 bp in length and 22 (2%) were greater than 100,000 bp in length. A bimodal distribution with peaks at ∼3000 bp and ∼70,000 bp was observed in both sets **(Fig. 1D)**. However, the distribution of the circular elements from the low [U] sites was skewed towards the smaller sizes (∼3000 bp peak). The average length of the circular elements from the high [U] samples was significantly longer (22,247 vs. 11,880 bp; two-tailed Welch’s t-test; *p = 0.0008*) than those from the low [U] samples. We next examined the structural and functional implications of this differential size distribution between these two groups of putative MGEs.

### Classification of circular elements

Expected circular MGEs in our dataset include plasmids and viruses as well as integrative and conjugative elements (ICEs), integrons, and transposons **(Fig. 2A, Table S6).** We identified 111 unique viral genomes (7% of all circular elements) with 39 from the high [U] sites and 72 from low [U] sites **(Fig. 2B, Table S7)**. The size distribution of viral assemblies was similar (two-tailed Welch’s t-test; *p > 0.05*) between the two datasets **(Fig. S1A)**. A similar proportion of predicted viral assemblies from high and low [U] sites clustered with known viral taxa within a gene-sharing network **(Fig. S1B, C)**. Lifestyle analysis predicted that, from the high [U] regions, 23/39 (59%) of the genomes were temperature phages while 15/39 (38%) of the genomes were virulent (*i.e.,* lytic) phages. While, from the low [U] regions, 20/72 genomes (28%) and 51/72 genomes (71%) were predicted temperate and virulent phages, respectively **(Fig. S1C)**

**Figure 2.**
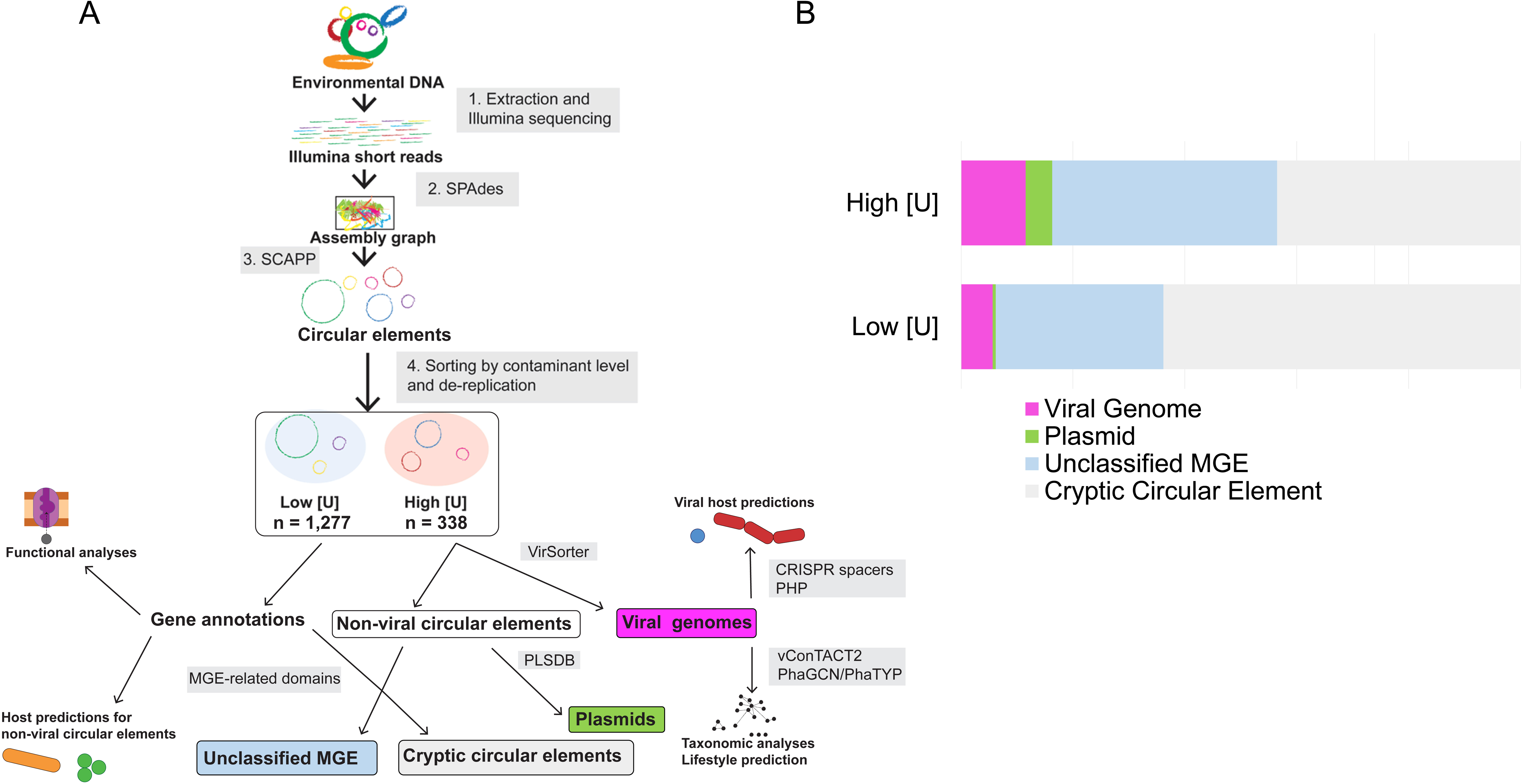
Classification of ORR MGEs. **(A)** Analysis workflow for the ORR mobilome. Illumina short reads were assembled using SPAdes and SCAPP was used to peel off putative circular mobile genetic elements (MGEs) from the SPAdes assembly graphs. Assemblies were then sorted by whether they originated from a high or low [U] sample and were de-replicated within those samples. Downstream analyses included MGE type classification, functional gene analysis, and host prediction. (B) Proportions of classified circular elements (left to right): pink indicates viral genomes that were predicted using VirSorter; green indicates assemblies predicted to be plasmids based off similarities to known plasmids in the PLSDB; light blue represents assemblies not classified by the prior two methods, but that carry MGE-related protein domains; and grey represents otherwise unknown (i.e., “cryptic”) circular elements.

We then compared the 1,504 non-viral circular elements to the PLSDB. Only 23 circular elements (16 from the high [U] sites and 7 from the low [U] sites) had similarity to known plasmids **(Fig. 2A, B, Table S8)**. These known plasmids were associated with various bacterial hosts and originated from both animal and environmental samples. The most common hosts were *Acinetobacter* and *Sphingobium* for the high and low [U] regions, respectively. The remaining unclassified circular elements were then analyzed for known MGE-related Pfam domains **(Table S4, Table S6)**^19^. From the high [U] sites, 121 circular elements were identified with MGE-related Pfams, including 75 (62%) with plasmid replication domains. From the low [U] sites, an additional 279 circular elements were identified, including 54 (19%) with plasmid replication domains **(Table S9)**. As highlighted with the plasmid replication domains, these “unclassified MGE” populations are distinct between the high and low [U] sites **(Figure S2).** We further annotated the remaining 1,104 circular elements using several additional protein domain databases (see methods section for complete list), allowing us to identify 15 and 104 additional circular elements with MGE-related domains from the high and low [U] regions, respectively **(Fig. 2A, B, Table S6).**

The remaining 43% and 64% of the circular elements from the high and low [U] sites, respectively, are cryptic, with no known domains involved in MGE replication or mobilization **(Fig. 2A, Table S6)**. While these cryptic circular elements were significantly shorter (AVG: 3,026 bp vs. 30,288 bp; two-tailed Welch’s t-test; *p = 4E-11*) **(Fig. S3)** than the non-cryptic elements, these sequences nonetheless encode genes that may serve important ecological functions **(Table S12)**. For example, the cryptic circular element AA_WF_A-C-Q_17, originating from high [U] groundwater, carries nine genes with putative metal resistance functions. While AA_WF_A-C-Q_17 does not encode any known MGE-related Pfams, a DUF6088 family protein is present on this assembly **(Table S10)**. This uncharacterized protein family is distantly related to the AbiEi antitoxin family of proteins, suggesting that this cryptic circular element could be a true novel MGE. Among the most common domains annotated on these cryptic circular elements included non-ribosomal peptide synthases involved in the biosynthesis of various secondary metabolites, chemotaxis proteins, pilus assembly proteins, and a variety of response regulators **(Table S12)**. Overall, we propose that the difference in circular element size distributions (**Fig. 1D)** between the two sample sets is attributable to the distinct populations of MGEs present.

### Predicted hosts of circular elements

Of the viral genomes from the high [U] regions, 47% were predicted by *k*-mer similarity to infect Proteobacteria hosts **(Fig. 3A, Table S7)** compared to 29% of the viral genomes from the low [U] regions. In the high [U] regions of the ORR, Proteobacteria predominate, while pristine regions are characterized by greater microbial diversity ^55,56^. Several Proteobacteria genera in the high [U] regions of the ORR are believed to play significant roles in nitrogen cycling in the subsurface, a process of critical interest due to the elevated levels of nitrate contaminating these areas of the ORR ^55^. For example, denitrifying *Rhodanobacter* spp. dominate in the most contaminated regions of the ORR subsurface. A previous study found that 82% of the microbial community in contaminated (*i.e.,* high [U]) groundwater well FW106 was *Rhodanobacter* ^56^.

**Figure 3.**
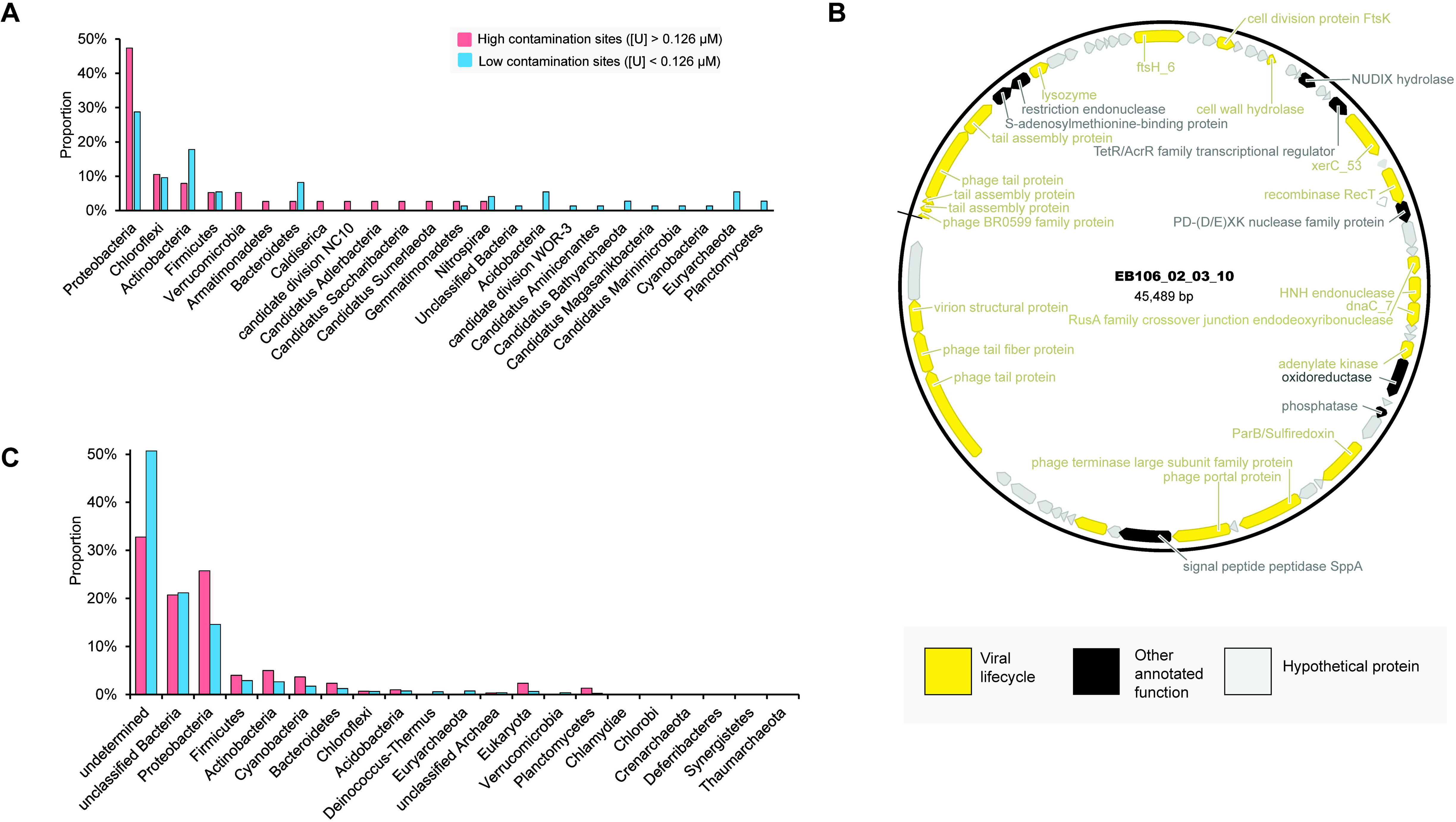
Prediction of circular element hosts. (A) Predictions of viral host phyla using a *k*-mer based method (PHP). Proportions were determined by normalizing against the total number of unique circular elements in each sample set. (B) Viral genome map with CRISPR-spacer-predicted *Rhodanobacter* host. (C) Predictions of host phyla for non-viral circular elements using a gene phylogeny-based approach. Proportions were determined by normalizing against the total number of unique circular elements in each sample set.

In parallel, we performed CRISPR spacer analysis against the IMG/VR database to determine more specific taxonomic assignments. Five out of 111 viral genomes were matched to an isolate host. An additional, 13 viral genomes were matched to metagenome-associated hosts. As an example, we identified a putative *Rhodanobacter* phage (EB106_02_03_10) **(Fig 3B)** in the high [U] samples. It is a predicted temperate phage and its genome encodes XerC and RecT-type recombinases that may facilitate integration in the host genome **(Table S10)**. Interestingly, the EB106_02_03_10 genome encodes a *Bgl*II endonuclease that may provide host immunity against simultaneous infection when this phage is in its lysogenic life stage ^57^. Also present in the genome is a gene encoding a putative oxidoreductase, but it is unclear if this represents a novel AMG or a phage lifecycle-related gene. This phage genome also encodes two different cell wall hydrolases that can promote cell lysis and drive biomass turnover in the ORR subsurface ^58–60^.

Host predictions for the remaining (*i.e.*, non-viral) circular elements were performed using gene taxonomy information extracted from the eggNOG-mapper annotation files **(Table S6)**. Due to the absence of known genes, a host could not be determined for 53% of the non-viral circular elements. Like the viral genomes, the most frequently identified host phylum in both sets was the Proteobacteria, with a greater proportion observed among the circular elements originating from high [U] samples **(Fig 3C)**. Where matches were previously made to plasmids in the PLSDB **(Table S8)**, host predictions were cross-checked with host metadata. Out of the 23 assemblies with PLSDB hits, 21 had predicted host taxonomy that was consistent with the host metadata of the PLSDB plasmid.

### Functional analysis of mobile genetic element assemblies

We next compared the functional gene content of the circular elements from high and low [U] regions of the ORR subsurface. We hypothesized that both heavy metal resistance genes (HMRGs) and antibiotic resistance genes (ARGs) would be enriched in the high [U] circular elements due to the selective pressure of the heavy metal contamination.

### Minimal enrichment of antibiotic resistance genes (ARGs) in high [U] sites

In some environments, heavy metal contamination can co-select for antibiotic resistance genes (ARGs) alongside HMRGs^61^ ^62,63^. However, the ARG count in our complete dataset was low **(Table S11).** The ARG counts were normalized against the total CDS count for each sample set (*i.e.,* high [U] and low [U]) to account for the size differences in the circular elements noted in the prior section. The overall ARG abundance was similar between the high and low [U] sample sets (Two-tailed Fisher’s exact test, *p > 0.05*) **(Fig. S4A)**. When considering individual genes, no enrichment pattern was observed, with three exceptions **(Fig. S5B)**. The ARG *acrA* (resistance to tetracyclines, penams, fluoroquinolones, rifamycins, and phenicols) is enriched in the high [U] samples (Two-tailed Fisher’s exact test, *p < 0.01*). AcrAB is a multidrug efflux protein with broad specify that has been implicated in the transport of other compounds related to metal homeostasis such as siderophores^64^. Previously, we found that iron homeostasis is significantly dysregulated in the multi-metal contaminated environments like the ORR subsurface ^65^. In contrast, *ampC* (resistance to penams and cephalosporins) and *sunT* (resistance to lantibiotics) are enriched in MGEs from low [U] samples. We repeated this analysis in a more conservative manner by first removing the cryptic circular elements from the dataset. This calculation yielded similar results to the analysis that had included the cryptic circular elements **(Table S13).** Overall, these data do not support a substantial enrichment of ARG among our putative MGEs from the high [U] samples.

### Assemblies from high [U] sites are enriched in heavy metal resistance gene clusters

As MGEs are frequently vectors of HMRGs^12,66^, we next examined our dataset for genes involved in heavy metal resistance **(Table S2, S10, S11)**. We found a significant (Two-tailed Fisher’s exact test, *p < 0.0001*) overabundance of HMRG content on the circular elements from the high [U] sites **(Fig. S5A)**. The assemblies from the high [U] regions encoded 32 HMRGs per 1 million base pairs (∼6.4% of total CDS) while the assemblies from low [U] sites encoded 6 HMRGs per 1 million base pairs (∼1.5% CDS). This trend was largely replicated in the analysis of individual HMRGs **(Fig. 4A).** Major exceptions included genes involved in arsenic (*acr3, arsB, arsC, arsA*) and tellurium resistance (*terC, tehB*) where no enrichment was observed in either direction. No HMRGs were enriched on the circular elements from the low [U] sites. We repeated this analysis using datasets that excluded the cryptic elements which yielded identical results to the previous analysis with the cryptic elements **(Table S14).**

**Figure 4.**
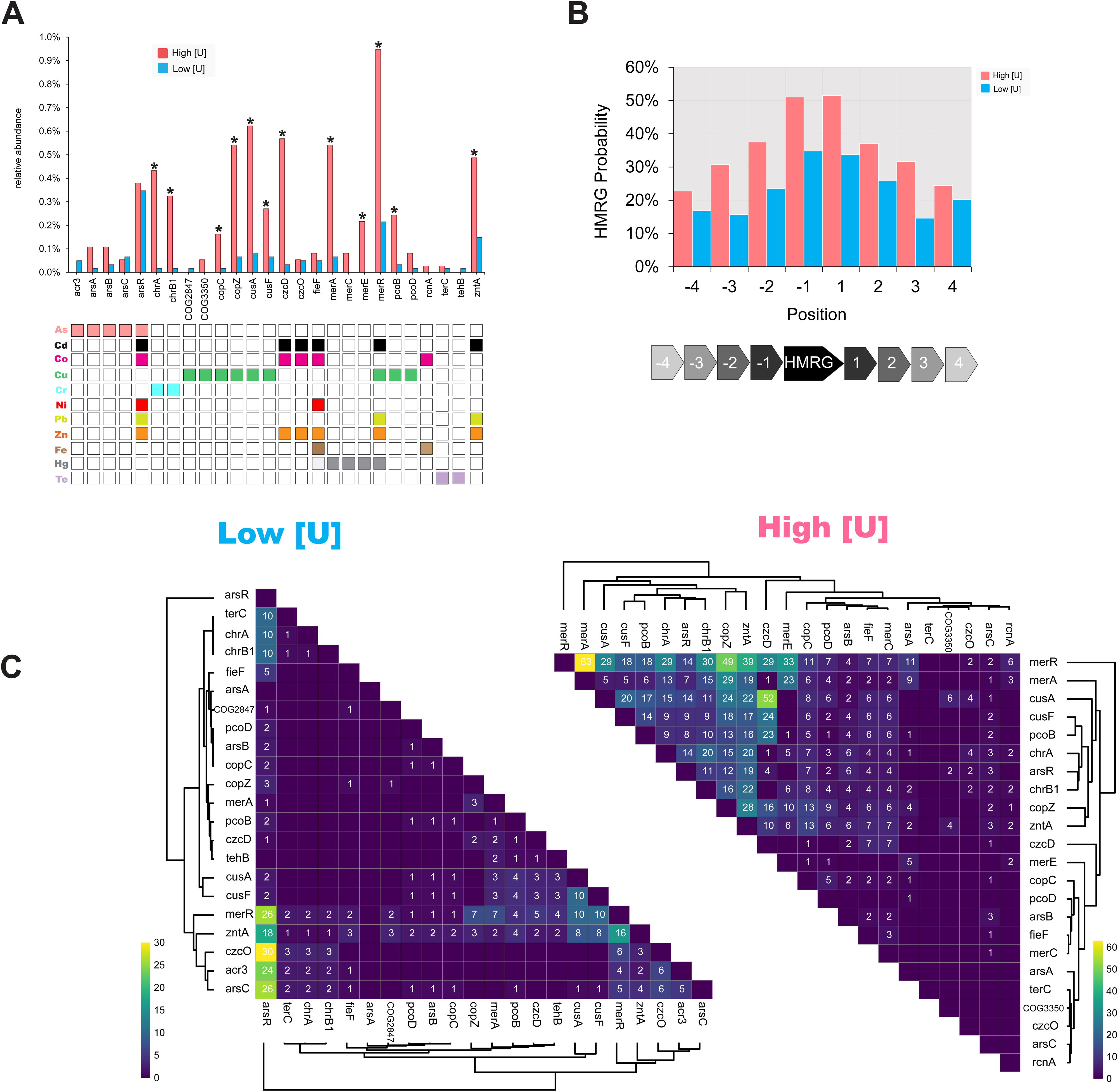
Analysis of MGE-associated HMRGs. (A) Relative abundance of individual HMRGs normalized against the total annotated CDS in each de-replicated dataset. Statistical comparisons were performed using a two-sided Fisher’s exact test (* p < 0.05; n.s = not significant). The associated grid indicates metals that each gene confers resistance to. (B) Neighborhood analysis of individual HMRGs. The histogram displays the probability that an individual HMRG is immediately adjacent (+/-4 ORFs) to another HMRG. (C) The heat map displays co-occurrence patterns of HMRG pairings on circular elements. Only have of each matrix is shown for simplicity. Co-occurrence > 0 are shown on the heat map. Clustering was performed using a Euclidian distance metric.

We predicted that MGEs from higher [U] regions may be more likely to encode multiple HMRGs than those from the low [U] regions due to the multi-metal components of the contaminant plume. We found that HMRG on the MGEs from the high [U] regions are more likely to co-occur in proximity in the form of gene clusters than those from the low [U] regions **(Fig. 4B)**. We next analyzed specific HMRG co-occurrence patterns on the circular elements **(Fig 4C)**. Among the high [U] circular elements, we observed frequent co-occurrences of *merR, chrB1, chrA, copZ*, *merA, arsR, cusA, cusF, pcoB, czcD*, and *zntA* with each other. The HMRGs confer resistance to a wide range of metals. An example of this HMRG-clustering phenomenon is seen on the plasmid EB106_03_01_3 from the high [U] region. This plasmid contains *merA, merR, zntA, czcD*, and *arsR* in close physical proximity to each other (**Fig. S5B**). In contrast, the low [U] circular elements primarily had co-occurrences of individual HMRGs and their corresponding metal-responsive transcriptional regulators, *merR* and *arsR* **(Fig. S5B).**

### Partitioning of HMRGs between classes of MGEs

We hypothesized that both plasmids and viral genomes would be significant vectors of HMRGs in the ORR subsurface. Viruses that infect bacteria or archaea may significantly augment host metabolic potential during infection via auxiliary metabolic genes (AMGs). AMGs encode diverse functions such as nutrient metabolism ^67^, virulence factors ^68^, oxidative stress defense ^69^, and toxicant (*e.g.*, antibiotics and heavy metals) resistance ^70^. We identified AMGs in our predicted viral assemblies by excluding genes involved in viral replication, structure, and function **(Table S15).** No AMGs related to heavy metal resistance were found in the assemblies from the low [U] regions. Within the high [U] regions, a single viral assembly (AA_WF_G-H-W_1) carried AMGs that may function in heavy metal resistance including TerD domain containing proteins and TelA family protein which have annotated functions in tellurium resistance but likely represent general cell envelope stress resistance proteins based on more recent research **(Table S15)**^71,72^. Additionally, a gene encoding a TerC family protein was present; however, TerC was recently found to be involved in metalation of exoenzymes during protein secretion^73^. While Proteobacteria predominate in the contaminated ORR subsurface^26^, AA_WF_G-H-W_1 is predicted to infect a member of the phylum Caldiserica. Thus, the phages in our dataset more are likely to impact Proteobacteria population dynamics in the contaminated ORR subsurface through biomass turnover rather than the introduction of novel adaptive genes via HGT.

In contrast to the viral genomes, assemblies with similarity to known plasmids in the PLSDB (*i.e.*, predicted plasmids) carried a substantial proportion of the HMRGs (19% of HMRG-encoding MGEs) despite representing only 1% of the total MGEs in our dataset **(Fig. 5)**. The remaining HMRGs were associated with (1) the “unclassifiable” circular elements that encode MGE-related protein domains (70% of HMRG-encoding MGEs) and (2) the cryptic circular elements (9%) **(Fig. S3)**. We examined the host predictions for these non-viral MGEs that carry HMRGs. At the class level, Beta/Gammaproteobacteria are the most common hosts for the HMRGs from the high [U] regions while Alphaproteobacteria are the most common hosts of these genes in the low [U] regions **(Fig. 6B)**.

**Figure 5.**
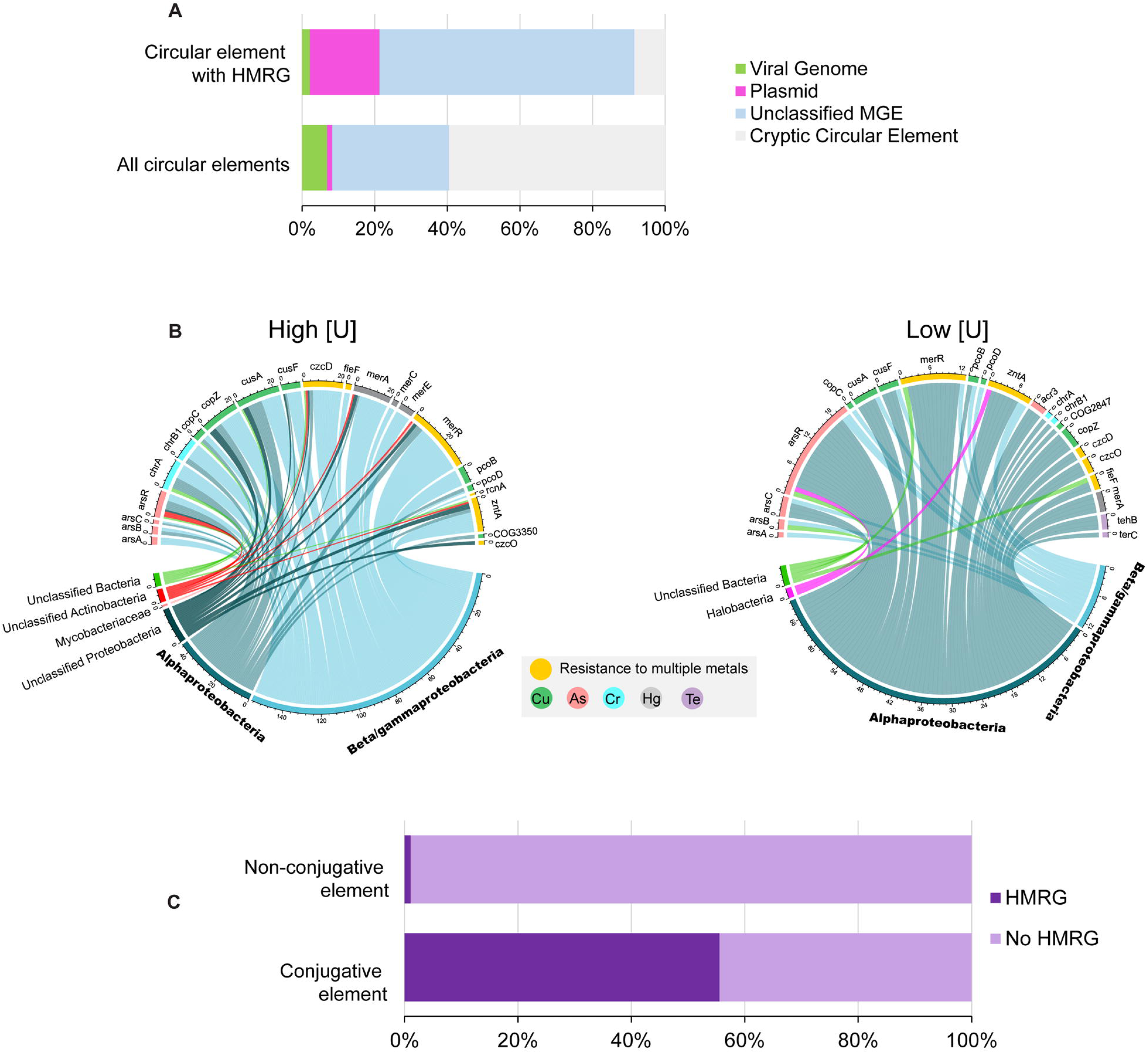
Partitioning of HMRGs between classes of MGEs and hosts. (A) The top bar shows the classifications of the circular elements that encode at least one HMRG (n = 47). The bottom bar shows the classifications of all circular elements in the dataset (n = 1,615). Proportions of classified circular elements (left to right): pink indicates viral genomes that were predicted using VirSorter; green indicates assemblies predicted to be plasmids based off similarities to known plasmids in the PLSDB; light blue represents assemblies not classified by the prior two methods, but that carry MGE-related protein domains; and grey represents otherwise unknown (i.e., “cryptic”) circular elements. (B) Chord diagrams linking HMRG (top of diagram) in the dataset to associated host class (bottom). (C) Proportions of conjugative and non-conjugative elements that carry 1+ HMRG (dark purple) or no HMRG (light purple). Proportions are normalized against the total number of unique assemblies. Viral genomes were excluded from this analysis.

We next explored the possible mechanisms by which the MGE-associated HMRGs could move within the microbial populations in the ORR subsurface. As described above, the viral genomes in our dataset are not the main vectors for the movement of HMRG within the ORR microbial populations. However, 54 of the non-viral circular elements carried at least one COG associated with conjugal transfer systems found on conjugative (*i.e.*, self-transmissible) plasmids and ICEs. We examined the co-occurrence patterns of HMRGs and genes encoding conjugative transfer machinery **(Table S3)**. We found that the HMRGs in our dataset were significantly more likely to be associated with a conjugative element than a non-conjugative element (two-tailed Fisher’s exact test, *p < 0.0001*) **(Fig. 5C)**. In fact, 90% of the identified HMRG were located on a conjugative element. These trends remained even when the data were analyzed by contamination levels (*i.e.,* high vs. low [U]). Thus, most HMRGs in our dataset have a high potential for future conjugative transfer within the ORR community independent of the current level of contamination at their origin site.

## DISCUSSION

This is the first study to characterize the mobilome of a highly contaminated subsurface environment. Through the usage of short-read sequences alone, we recovered 1,615 unique circular elements that we propose represent MGEs. In this dataset, we simultaneously identified and characterized viral genomes, plasmids, and other types of MGEs, including potentially novel families. Thus, this work diverges from prior studies that have primarily focused on one class of MGEs (*e.g.,* plasmids, viruses) within the mobilome. This agnostic approach allowed for examination of partitioning of key functional genes between different classes of MGEs. For example, we found that the HMRGs in our dataset were exclusively associated with non-viral contigs—many of which were predicted conjugative elements. Additionally, we speculate that the lack of a “plasmid safe” DNA digestion step employed by previous studies allowed us to recover a larger number of substantially longer putative MGEs than prior mobilome studies^19^.

Environmental stressors may select for a pool of mobile genetic elements that confer fitness advantages to their hosts within the specific environmental context^13,19,74,75^. We observed an enrichment of multiple HMRGs in our mobilome extracted from samples from high [U] sites. The HMRG enrichment pattern was consistent with the contaminant profile of the ORR subsurface. For example, we observe enrichment of genes involved in Cd, Cu, and Co resistance—all of which are found at sufficiently elevated concentrations in the contaminated ORR subsurface to be toxic to certain native microbiota^76–78^. In contrast, abundance of genes involved in Te and As resistance were similar between the two datasets. In the contaminated regions of the ORR, arsenic is slightly elevated relative to the background ([As]_Median_ = 5 nM vs. 0.5 nM); however, these values are orders of magnitude below typical toxicity thresholds for arsenic in microorganisms^76^. Likewise, tellurium is an exceptionally rare element that is not a component of the ORR contamination plume ^26^. Additionally, several studies have also suggested that the classical “tellurium resistance genes” (*e.g., ter, teh*) may have primary functions unrelated to tellurium resistance, such as phage and antibiotic resistance ^79,80^.

A distinctive feature of the MGE-associated HMRGs from the high [U] regions of the ORR was their physical clustering on the circular elements. The non-random associations of functionally similar genes are well-characterized in bacteria^81^. However, the evolutionary mechanisms driving this clustering remain controversial and are likely to be context specific. The Selfish Operon Model proposes that gene clusters are formed and maintained through the emergent benefit to the genes themselves rather than their host organism ^81^. In this model, the clustering of genes is essential for the successful horizontal transfer of genes that individually confer minimal selectable function. For example, clustering of multiple antibiotic resistance genes on plasmids can be explained by an extension of the selfish operon theory^3^. Co-localization of antibiotic resistance genes on a single plasmid may be a successful strategy for long-term antibiotic resistance gene persistence in human pathogens that are frequently targeted by multi-antibiotic therapy. We can apply a similar line of logic to the HMRGs in our dataset: individual HMRGs may provide minimal selectable function in a complex high-stress environment, such as the ORR subsurface, where multiple metal contaminants co-exist. Successfully retained HMRGs may be frequently co-localized with other HMRGs. In a larger-scale example of this phenomenon, Staehlin, et al. ^82^ used a molecular clock to link the origin and diversification of a 19 gene copper homeostasis and silver resistance transposon in *Enterobacteriaceae* to increases in human metallurgical activity throughout history.

In our study, a substantial proportion of the MGE-encoded HMRGs (all sites: 94%; H: 94%; L: 93%) were associated with Proteobacteria hosts. A recent study by Finks and Martiny ^83^ found that plasmid traits varied significantly with their host’s taxonomic assignment at the phylum level using a large dataset of publicly available plasmids. However, certain trait variation was still controlled, in part, by differences in environment of origin. When we only consider circular elements with predicted Proteobacteria hosts, the HMRG content remains significantly enriched (two-tailed Fisher’s exact test, *p < 0.0001)* among those originating from high [U] regions of the ORR subsurface compared to those from the low [U] regions. Considering our findings here in the context of prior work^83,84^, we propose a model for our system where host taxonomy establishes a baseline “genetic potential” for acquisition of novel MGE-encoded genes. This genetic potential may be controlled, in part, by positive or negative interactions between horizontally acquired resistance genes and cellular metabolic genes in the host genome^85^. However, selective pressures in the host’s environment control the enrichment of certain MGE-encoded traits within a taxonomic group. As a final point of consideration: HMRG are best characterized in readily culturable Proteobacteria^86^. Thus, the association between the two in our environment may reflect, to an extent, this under-sampling in the scientific literature. However, this association is not entirely spurious as Proteobacteria are indeed highly enriched within the high [U] ORR subsurface relative to low [U] regions of the site^55^. The ability to rapidly acquire and maintain MGEs carrying clusters of HMRGs may have been one factor that has facilitated the success of members of this phylum in the ORR subsurface.

Viral genomes have been proposed as significant vectors for the shuttling of metabolic genes between different host cells^67^. However, we identified only a singular instance of a HMRG encoded on a viral genome. Instead, we found that the HMRGs in our dataset were largely associated with conjugative elements. These “conjugative elements” likely include both conjugative plasmids and integrative and conjugative elements^87^. These HMRG-carrying conjugative elements included both (1) circular elements high similarity to known plasmids in the PLSDB as well as (2) potentially novel conjugative elements. Our results mirror those of Finks and Martiny ^83^ who found a significant association between resistance genes (*i.e.,* antibiotic, metal, and biocide resistance) and the MOB family relaxase which is found on both conjugative and mobilizable plasmids. The underlying mechanism for this association between resistance genes and conjugation/mobilization genes is unclear and warrants further investigation. One possibility is that simple genetic traits conferring a strong and immediate selective advantage (*i.e*., resistance genes) may increase the fitness of the new host strain, offsetting the fitness costs of maintaining these larger plasmids during the initial period of plasmid-host adaptation following acquisition, promoting the linkage of resistance genes to these classes of larger plasmids^88^.

Consistent with other metagenomic studies, we have uncovered an enormous diversity of potentially novel MGEs in the ORR subsurface. Across our entire dataset, 60% of the circular DNA elements did not encode any identifiable MGE-related genes. This is similar to the results of Kirstahler, et al. ^19^ who found that 53% of the circular DNA elements in their global sewage mobilome did not carry any identifiable MGE-related Pfam domains. However, several studies have identified small cryptic plasmids in bacterial isolates that contain no recognizable replication machinery^89–92^. It has been speculated that cryptic plasmids may play roles in viral defense^93^ or functionally serving as empty backbones acquisition of novel genes^94^ Additionally, some of these cryptic elements may represent currently undescribed classes of MGEs. In recent years, numerous new classes of MGEs have been identified and described in the literature^95,96^. We expect that as time goes on, our ability to better classify these currently cryptic elements will improve. One limitation of this work is our usage of short-read sequencing technology, increasing the likelihood that some of these cryptic elements may represent misassembles due to failure to span repetitive elements of larger sequences^97^. However, we did remove the obvious instances of this occurrence from our analysis. Nonetheless, our work here highlights the value of assembling completed mobile genetic elements from metagenomic datasets. These data provide useful genomic and taxonomic context to key resistance traits often considered in isolation in metagenomic studies.

## DATA AVAILABILITY

Accession numbers for metagenome raw reads are listed in **Table S5**. The sequences of the SCAPP-generated circular elements are available in a public KBase narrative: https://doi.org/10.25982/160429.7/2203457 [for review purposes: https://kbase.us/n/160429/7/]

## ACKNOWLEDGEMENTS

This material by ENIGMA (Ecosystems and Networks Integrated with Genes and Molecular Assemblies) (http://enigma.lbl.gov), a Science Focus Area Program at Lawrence Berkeley National Laboratory, is based on work supported by the U.S. Department of Energy, Office of Science, Office of Biological and Environmental Research, under contract DE-AC02-05CH11231.

## REFERENCES

1 Siefert, J. L. Defining the mobilome. Methods Mol Biol 532, 13–27 (2009). 10.1007/978-1-60327-853-9_2

2 van Elsas, J. D. & Bailey, M. J. The ecology of transfer of mobile genetic elements. FEMS Microbiol. Ecol. 42, 187–197 (2002). 10.1111/j.1574-6941.2002.tb01008.x

3 Wiedenbeck, J. & Cohan, F. M. Origins of bacterial diversity through horizontal genetic transfer and adaptation to new ecological niches. FEMS Microbiology Reviews 35, 957–976 (2011). 10.1111/j.1574-6976.2011.00292.x

4 Bhatt, P. et al. Plasmid-mediated catabolism for the removal of xenobiotics from the environment. J. Hazard. Mater. 420, 126618 (2021).

5 Bennett, P. M. Plasmid encoded antibiotic resistance: acquisition and transfer of antibiotic resistance genes in bacteria. British journal of pharmacology 153, S347–S357 (2008).

6 Li, J. et al. Toxin plasmids of Clostridium perfringens. Microbiol. Mol. Biol. Rev. 77, 208–233 (2013).

7 Bruins, M. R., Kapil, S. & Oehme, F. W. Microbial Resistance to Metals in the Environment. Ecotoxicol. Environ. Saf. 45, 198–207 (2000). 10.1006/eesa.1999.1860

8 Gillings, M. R. Lateral gene transfer, bacterial genome evolution, and the Anthropocene. Annals of the New York Academy of Sciences 1389, 20–36 (2017). 10.1111/nyas.13213

9 Brooks, S. C. Waste characteristics of the former S-3 ponds and outline of uranium chemistry relevant to NABIR Field Research Center studies. (NABIR Field Research Center, Oak Ridge, Tenn, 2001).

10 Hemme, C. L. et al. Lateral gene transfer in a heavy metal-contaminated-groundwater microbial community. mBio 7, e02234–02215 (2016). doi:10.1128/mBio.02234-15

11 Peng, M. et al. Genomic Features and Pervasive Negative Selection in *Rhodanobacter* Strains Isolated from Nitrate and Heavy Metal Contaminated Aquifer. Microbiology Spectrum 10, e02591–02521 (2022). doi:10.1128/spectrum.02591-21

12 Martinez, R. J. et al. Horizontal gene transfer of PIB-type ATPases among bacteria isolated from radionuclide-and metal-contaminated subsurface soils. Appl Environ Microbiol 72, 3111–3118 (2006). 10.1128/aem.72.5.3111-3118.2006

13 Perez, M. F. et al. First Report on the Plasmidome From a High-Altitude Lake of the Andean Puna. Front. Microbiol. 11 (2020). 10.3389/fmicb.2020.01343

14 Kav, A. B. et al. Insights into the bovine rumen plasmidome. Proc. Natl. Acad. Sci. U. S. A. 109, 5452–5457 (2012).

15 Perez, M. F. et al. Assessment of the plasmidome of an extremophilic microbial community from the Diamante Lake, Argentina. Sci. Rep. 11, 21459 (2021).

16 Gulino, K. et al. Initial mapping of the New York City wastewater virome. Msystems 5, e00876–00819 (2020).

17 Trubl, G. et al. Soil Viruses Are Underexplored Players in Ecosystem Carbon Processing. mSystems 3, e00076–00018 (2018). doi:10.1128/mSystems.00076-18

18 Kothari, A. et al. Large Circular Plasmids from Groundwater Plasmidomes Span Multiple Incompatibility Groups and Are Enriched in Multimetal Resistance Genes. mBio 10, e02899–02818 (2019). 10.1128/mBio.02899-18

19 Kirstahler, P., Teudt, F., Otani, S., Aarestrup, F. M. & Pamp, S. J. A Peek into the Plasmidome of Global Sewage. mSystems 6, e00283–00221 (2021). doi:10.1128/mSystems.00283-21

20 Hall, J. P., Botelho, J., Cazares, A. & Baltrus, D. A. What makes a megaplasmid? Philosophical Transactions of the Royal Society B 377, 20200472 (2022).

21 Zrelovs, N., Dislers, A. & Kazaks, A. Motley Crew: Overview of the Currently Available Phage Diversity. Front. Microbiol. 11 (2021). 10.3389/fmicb.2020.579452

22 Pellow, D. et al. SCAPP: an algorithm for improved plasmid assembly in metagenomes. Microbiome 9, 144 (2021). 10.1186/s40168-021-01068-z

23 Lui, L. M., Nielsen, T. N. & Arkin, A. P. A method for achieving complete microbial genomes and improving bins from metagenomics data. PLoS Comput. Biol. 17, e1008972 (2021). 10.1371/journal.pcbi.1008972

24 Lui, L. M. et al. Mechanism across scales: a holistic modeling framework integrating laboratory and field studies for microbial ecology. Front. Microbiol. 12, 642422 (2021). 10.3389/fmicb.2021.642422

25 Khedkar, S. et al. Landscape of mobile genetic elements and their antibiotic resistance cargo in prokaryotic genomes. Nucleic Acids Res. 50, 3155–3168 (2022). 10.1093/nar/gkac163

26 Smith, M. B. et al. Natural bacterial communities serve as quantitative geochemical biosensors. mBio 6, e00326–00315 (2015). doi:10.1128/mBio.00326-15

27 Wilpiszeski, R. L. et al. In-field bioreactors demonstrate dynamic shifts in microbial communities in response to geochemical perturbations. PLoS One 15, e0232437 (2020). 10.1371/journal.pone.0232437

28 Gushgari-Doyle, S. et al. Genotype to ecotype in niche environments: adaptation of Arthrobacter to carbon availability and environmental conditions. ISME Communications 2, 32 (2022). 10.1038/s43705-022-00113-8

29 Watson, D., Kostka, J., Fields, M. & Jardine, P. The Oak Ridge field research center conceptual model. NABIR Field Research Center, Oak Ridge, TN (2004).

30 Thorgersen, M. P. et al. Molybdenum availability Is key to nitrate removal in contaminated groundwater environments. Applied and Environmental Microbiology 81, 4976–4983 (2015). doi:10.1128/AEM.00917-15

31 Cantalapiedra, C. P., Hernández-Plaza, A., Letunic, I., Bork, P. & Huerta-Cepas, J. eggNOG-mapper v2: Functional Annotation, Orthology Assignments, and Domain Prediction at the Metagenomic Scale. Mol. Biol. Evol. 38, 5825–5829 (2021). 10.1093/molbev/msab293

32 Galperin, M. Y. et al. COG database update: focus on microbial diversity, model organisms, and widespread pathogens. Nucleic Acids Res. 49, D274–D281 (2021). 10.1093/nar/gkaa1018

33 Marchler-Bauer, A. et al. CDD/SPARCLE: functional classification of proteins via subfamily domain architectures. Nucleic Acids Res. 45, D200–D203 (2017). 10.1093/nar/gkw1129

34 Schultz, J., Copley, R. R., Doerks, T., Ponting, C. P. & Bork, P. SMART: a web-based tool for the study of genetically mobile domains. Nucleic Acids Res. 28, 231–234 (2000). 10.1093/nar/28.1.231

35 Klimke, W. et al. The National Center for Biotechnology Information’s Protein Clusters Database. Nucleic Acids Res. 37, D216–D223 (2009). 10.1093/nar/gkn734

36 Haft, D. H. et al. TIGRFAMs and Genome Properties in 2013. Nucleic Acids Res. 41, D387–D395 (2013). 10.1093/nar/gks1234

37 Tatusova, T. et al. NCBI prokaryotic genome annotation pipeline. Nucleic Acids Res. 44, 6614–6624 (2016). 10.1093/nar/gkw569

38 Roux, S., Enault, F., Hurwitz, B. L. & Sullivan, M. B. VirSorter: mining viral signal from microbial genomic data. PeerJ 3, e985 (2015).

39 Arkin, A. P. et al. KBase: The United States Department of Energy Systems Biology Knowledgebase. Nat. Biotechnol. 36, 566–569 (2018). 10.1038/nbt.4163

40 Bin Jang, H., et al. Taxonomic assignment of uncultivated prokaryotic virus genomes is enabled by gene-sharing networks. Nat. Biotechnol. 37, 632–639 (2019). 10.1038/s41587-019-0100-8

41 Lefkowitz, E. J. et al. Virus taxonomy: the database of the International Committee on Taxonomy of Viruses (ICTV). Nucleic acids research 46, D708–D717 (2018).

42 Shang, J., Jiang, J. & Sun, Y. Bacteriophage classification for assembled contigs using graph convolutional network. Bioinformatics 37, i25–i33 (2021). 10.1093/bioinformatics/btab293

43 Shang, J., Tang, X. & Sun, Y. PhaTYP: predicting the lifestyle for bacteriophages using BERT. Brief. Bioinform. 24, bbac487 (2023). 10.1093/bib/bbac487

44 Finn, R. D. et al. The Pfam protein families database. Nucleic Acids Res. 38, D211–D222 (2009). 10.1093/nar/gkp985

45 Lu, C. et al. Prokaryotic virus host predictor: a Gaussian model for host prediction of prokaryotic viruses in metagenomics. BMC Biol. 19, 5 (2021). 10.1186/s12915-020-00938-6

46 Camargo, A. P. et al. IMG/VR v4: an expanded database of uncultivated virus genomes within a framework of extensive functional, taxonomic, and ecological metadata. Nucleic Acids Res. 51, D733–D743 (2023). 10.1093/nar/gkac1037

47 Gu, Z., Gu, L., Eils, R., Schlesner, M. & Brors, B. circlize implements and enhances circular visualization in R. Bioinformatics 30, 2811–2812 (2014). 10.1093/bioinformatics/btu393

48 Kolde, R. & Kolde, M. R. Package ‘pheatmap’. R package 1, 790 (2015).

49 Wickham, H., Chang, W. & Wickham, M. H. Package ‘ggplot2’. Create elegant data visualisations using the grammar of graphics. Version 2, 1–189 (2016).

50 Shannon, P. et al. Cytoscape: a software environment for integrated models of biomolecular interaction networks. Genome Res. 13, 2498–2504 (2003).

51 Tian, R. et al. Small and mighty: adaptation of superphylum Patescibacteria to groundwater environment drives their genome simplicity. Microbiome 8, 51 (2020). 10.1186/s40168-020-00825-w

52 Wu, X. et al. Distinct Depth-Discrete Profiles of Microbial Communities and Geochemical Insights in the Subsurface Critical Zone. Applied and Environmental Microbiology 89, e00500–00523 (2023). 10.1128/aem.00500-23

53 Lui Lauren, M., et al. Sediment and Groundwater Metagenomes from Subsurface Microbial Communities from the Oak Ridge National Laboratory Field Research Center, Oak Ridge, TN, USA. Microbiology Resource Announcements (pending).

54. Goff, J. L., et al. Supplemental Data for “Mixed waste contamination selects for a mobile genetic element population enriched in multiple heavy metal resistance genes”, <10.25982/160429.7/2203457> (2023).

55 Hemme, C. L. et al. Metagenomic insights into evolution of a heavy metal-contaminated groundwater microbial community. ISME J. 4, 660–672 (2010). 10.1038/ismej.2009.154

56 Hemme, C. L. et al. Comparative metagenomics reveals impact of contaminants on groundwater microbiomes. Front. Microbiol. 6 (2015). 10.3389/fmicb.2015.01205

57 Hampton, H. G., Watson, B. N. J. & Fineran, P. C. The arms race between bacteria and their phage foes. Nature 577, 327–336 (2020). 10.1038/s41586-019-1894-8

58 Daly, R. A. et al. Viruses control dominant bacteria colonizing the terrestrial deep biosphere after hydraulic fracturing. Nature Microbiology 4, 352–361 (2019). 10.1038/s41564-018-0312-6

59 Kuzyakov, Y. & Mason-Jones, K. Viruses in soil: Nano-scale undead drivers of microbial life, biogeochemical turnover and ecosystem functions. Soil Biology and Biochemistry 127, 305–317 (2018). 10.1016/j.soilbio.2018.09.032

60 Starr, E. P. et al. Stable-isotope-informed, genome-resolved metagenomics uncovers potential cross-kingdom interactions in rhizosphere soil. Msphere 6, e00085–00021 (2021).

61 Baker-Austin, C., Wright, M. S., Stepanauskas, R. & McArthur, J. V. Co-selection of antibiotic and metal resistance. Trends in Microbiology 14, 176–182 (2006). 10.1016/j.tim.2006.02.006

62 Liu, J.-l., et al. Metagenomic exploration of multi-resistance genes linked to microbial attributes in active nonferrous metal(loid) tailings. Environmental Pollution 273, 115667 (2021). 10.1016/j.envpol.2020.115667

63 Zou, H.-Y. et al. Antibiotic resistance genes in surface water and groundwater from mining affected environments. Science of The Total Environment 772, 145516 (2021). 10.1016/j.scitotenv.2021.145516

64 Horiyama, T. & Nishino, K. AcrB, AcrD, and MdtABC multidrug efflux systems are involved in enterobactin export in Escherichia coli. PLoS One 9, e108642 (2014).

65 Goff, J. L. et al. Mixed heavy metal stress induces global iron starvation response. The ISME Journal (2022). 10.1038/s41396-022-01351-3

66 Coombs, J. M. & Barkay, T. Molecular evidence for the evolution of metal homeostasis genes by lateral gene transfer in bacteria from the deep terrestrial subsurface. Applied and environmental microbiology 70, 1698–1707 (2004). 10.1128/AEM.70.3.1698-1707.2004

67 Luo, X.-Q. et al. Viral community-wide auxiliary metabolic genes differ by lifestyles, habitats, and hosts. Microbiome 10, 190 (2022). 10.1186/s40168-022-01384-y

68 Boyd, E. F. Bacteriophage-encoded bacterial virulence factors and phage–pathogenicity island interactions. Adv. Virus Res. 82, 91–118 (2012).

69 De Smet, J. et al. High coverage metabolomics analysis reveals phage-specific alterations to Pseudomonas aeruginosa physiology during infection. The ISME Journal 10, 1823–1835 (2016). 10.1038/ismej.2016.3

70 Hargreaves, K. R., Kropinski, A. M. & Clokie, M. R. Bacteriophage behavioral ecology: how phages alter their bacterial host’s habits. Bacteriophage 4, e85131 (2014).

71 Anantharaman, V., Iyer, L. M. & Aravind, L. Ter-dependent stress response systems: novel pathways related to metal sensing, production of a nucleoside-like metabolite, and DNA-processing. Mol. Biosyst. 8, 3142–3165 (2012). 10.1039/C2MB25239B

72 Collins, B., Joyce, S., Hill, C., Cotter, P. D. & Ross, R. P. TelA contributes to the innate resistance of Listeria monocytogenes to nisin and other cell wall-acting antibiotics. Antimicrob Agents Chemother 54, 4658–4663 (2010). 10.1128/aac.00290-10

73 He, B., Sachla, A. J. & Helmann, J. D. TerC proteins function during protein secretion to metalate exoenzymes. Nature Communications 14, 6186 (2023). 10.1038/s41467-023-41896-1

74 Zhang, T., Zhang, X.-X. & Ye, L. Plasmid Metagenome Reveals High Levels of Antibiotic Resistance Genes and Mobile Genetic Elements in Activated Sludge. PLoS One 6, e26041 (2011). 10.1371/journal.pone.0026041

75 Ji, M. et al. Tundra Soil Viruses Mediate Responses of Microbial Communities to Climate Warming. mBio 0, e03009–03022 doi:10.1128/mbio.03009-22

76 Carlson, H. K. et al. The selective pressures on the microbial community in a metal-contaminated aquifer. ISME J. 13, 937–949 (2019). 10.1038/s41396-018-0328-1

77 Goff, J. L. et al. Ecophysiological and genomic analyses of a representative isolate of highly abundant *Bacillus cereus* strains in contaminated subsurface sediments. Environmental Microbiology 24, 5546–5560 (2022). 10.1111/1462-2920.16173

78 Ge, X. et al. Characterization of a Metal-Resistant Bacillus Strain With a High Molybdate Affinity ModA From Contaminated Sediments at the Oak Ridge Reservation. Front. Microbiol. 11 (2020). 10.3389/fmicb.2020.587127

79 Whelan, K. F., Colleran, E. & Taylor, D. E. Phage inhibition, colicin resistance, and tellurite resistance are encoded by a single cluster of genes on the IncHI2 plasmid R478. J. Bacteriol. 177, 5016–5027 (1995). doi:10.1128/jb.177.17.5016-5027.1995

80 Turner, R. J., Taylor, D. E. & Weiner, J. H. Expression of Escherichia coli TehA gives resistance to antiseptics and disinfectants similar to that conferred by multidrug resistance efflux pumps. Antimicrobial agents and chemotherapy 41, 440–444 (1997).

81 Lawrence, J. Selfish operons: the evolutionary impact of gene clustering in prokaryotes and eukaryotes. Curr. Opin. Genet. Dev. 9, 642–648 (1999). 10.1016/S0959-437X(99)00025-8

82 Staehlin, B. M., Gibbons, J. G., Rokas, A., O’Halloran, T. V. & Slot, J. C. Evolution of a Heavy Metal Homeostasis/Resistance Island Reflects Increasing Copper Stress in Enterobacteria. Genome Biol. Evol. 8, 811–826 (2016). 10.1093/gbe/evw031

83 Finks, S. S. & Martiny, J. B. Plasmid-Encoded Traits Vary across Environments. Mbio, e03191–03122 (2023).

84 Redondo-Salvo, S. et al. Pathways for horizontal gene transfer in bacteria revealed by a global map of their plasmids. Nat. Commun. 11, 3602 (2020). 10.1038/s41467-020-17278-2

85 Palomino, A. et al. Metabolic genes on conjugative plasmids are highly prevalent in Escherichia coli and can protect against antibiotic treatment. The ISME Journal 17, 151–162 (2023). 10.1038/s41396-022-01329-1

86 Gillan, D. C. Metal resistance systems in cultivated bacteria: are they found in complex communities? Curr. Opin. Biotechnol. 38, 123–130 (2016). 10.1016/j.copbio.2016.01.012

87 Johnson, C. M. & Grossman, A. D. Integrative and Conjugative Elements (ICEs): What They Do and How They Work. Annu. Rev. Genet. 49, 577–601 (2015). 10.1146/annurev-genet-112414-055018

88 Prensky, H., Gomez-Simmonds, A., Uhlemann, A. C. & Lopatkin, A. J. Conjugation dynamics depend on both the plasmid acquisition cost and the fitness cost. Molecular systems biology 17, e9913 (2021).

89 Daming, R. et al. Complete DNA sequence and analysis of two cryptic plasmids isolated from Lactobacillus plantarum. Plasmid 50, 70–73 (2003). 10.1016/S0147-619X(03)00010-6

90 Prity, F. T. et al. The evolutionary tale of eight novel plasmids in a colistin-resistant environmental Acinetobacter baumannii isolate. Microb Genom 9 (2023). 10.1099/mgen.0.001010

91 Barry, K. E. et al. Don’t overlook the little guy: An evaluation of the frequency of small plasmids co-conjugating with larger carbapenemase gene containing plasmids. Plasmid 103, 1–8 (2019). 10.1016/j.plasmid.2019.03.005

92 Challacombe, J. F., Pillai, S. & Kuske, C. R. Shared features of cryptic plasmids from environmental and pathogenic Francisella species. PLOS ONE 12, e0183554 (2017). 10.1371/journal.pone.0183554

93 Ma, Y., Paulsen, I. T. & Palenik, B. Analysis of two marine metagenomes reveals the diversity of plasmids in oceanic environments. Environmental Microbiology 14, 453–466 (2012). 10.1111/j.1462-2920.2011.02633.x

94 Pérez-García, D. et al. Frequency and diversity of small plasmids in mesophilic Aeromonas isolates from fish, water and sediment. Plasmid 118, 102607 (2021). 10.1016/j.plasmid.2021.102607

95 Al-Shayeb, B. et al. Borgs are giant genetic elements with potential to expand metabolic capacity. Nature 610, 731–736 (2022). 10.1038/s41586-022-05256-1

96 Krupovic, M., Makarova, K. S., Forterre, P., Prangishvili, D. & Koonin, E. V. Casposons: a new superfamily of self-synthesizing DNA transposons at the origin of prokaryotic CRISPR-Cas immunity. BMC Biol. 12, 1–12 (2014).

97 Logsdon, G. A., Vollger, M. R. & Eichler, E. E. Long-read human genome sequencing and its applications. Nature Reviews Genetics 21, 597–614 (2020). 10.1038/s41576-020-0236-x

