## Supplemental Materials for "Mixed Waste Contamination Selects for a Mobile Genetic Element Population Enriched in Multiple Heavy Metal Resistance Genes"

**CONTENTS**

Tables S1-S12, S15…………………………………………………………………………...Excel files

Supplemental Methods…………………………………………………………………………………..2

Figure S1. Viral Genome Characteristics………………………………………………………………3

Figure S2. MGE-related Pfams on unclassifiable circular elements……………………………...4-5

Figure S3. Length distributions of cryptic vs. non-cryptic MGEs…………………………………….5

Figure S4. Analysis of MGE-associated antibiotic resistance genes (ARGs)……………………...6 Table S13. Abundance of ARG among non-cryptic circular elements……………………………...7

Figure S5. Analysis of MGE HMRG content…………………………………………………………..8

Table S14. Abundance of HMRG among non-cryptic circular elements……………………………9

**SUPPLEMENTAL METHODS**

***Metals analysis.*** For mercury analysis, the unfiltered groundwater samples collected into acid-washed polypropylene tubes (Sarstedt, Nümbrecht, Germany) and allowed to settle for few minutes. Then a subsample was diluted 1:10 into 0.5% (vol/wt) trace-grade HCl for mercury and 2% (vol/wt) trace-grade HNO_3_ for other metals (VWR, Radnor, PA) in acid-washed polypropylene tubes and stored up to a month at 4°C. The diluted, acidified, samples were spun down prior to metal analysis. The mercury samples were run separately due to the HCl preservation matrix, which was modified, from Louie et al. ^29^. The other metals including Cr, Mn, Fe, Co, Ni, Cu and Zn were analyzed based on EPA Method 6020A using an Agilent 7900 Inductively Coupled Plasma Mass Spectrometer (ICP-MS) fitted with MicroMist nebulizer, UHMI-spray chamber, Pt cones and an Octopole Reaction System (ORS) collision cell (Agilent Technologies, Santa Clara, CA). The data were collected using 3-point peak pattern, 3 replicates, 100 sweeps/rep and various integration times (all >0.3 s) using either no gas or He gas in the ORS collision cell. The external calibration standard for mercury, MSHG-10PPM (Inorganic Ventures, Christiansburg, Virginia), was used to create an 8-point curve from 0 - 10 ppb in 0.5% (vol/wt) trace-grade HCl (VWR, Radnor, PA). For all other metals the external calibration standards, IV-ICPMS-71A and IV-ICPMS-71B (Inorganic Ventures, Christiansburg, Virginia), were used to create a 12-point curve from 0 - 1000 pb for each element in 2% (vol/wt) trace-grade nitric acid (VWR, Radnor, PA). Both runs used the internal calibration standard, IV-ICPMS-71D (Inorganic Ventures, Christiansburg, Virginia), as it was continuously added to each sample to account for matrix effects. Data was processed using the nearest internal standard by mass and fitted to a linear curve though the calibration blank.

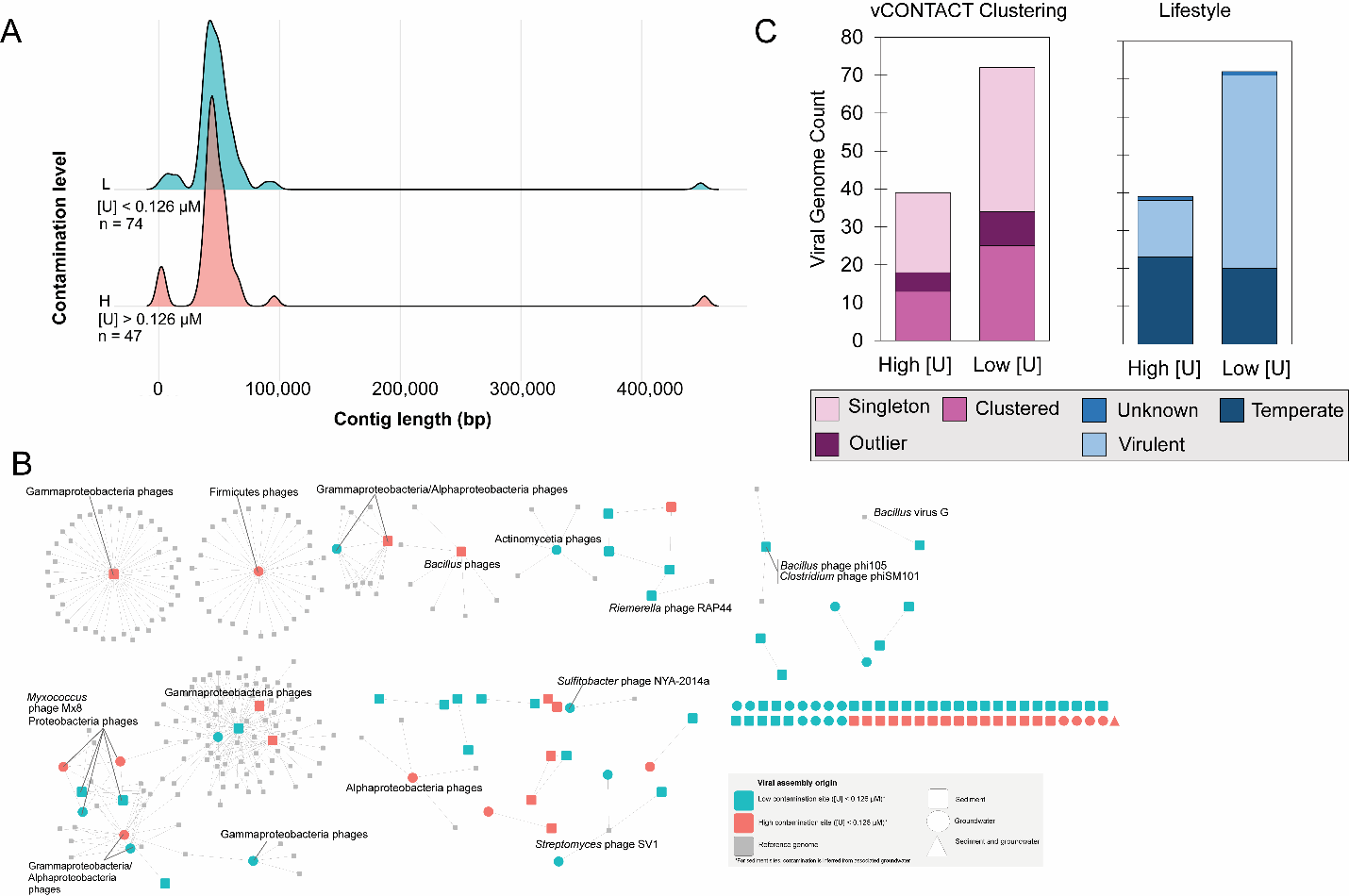

**Figure S1. Viral Genome Characteristics** (A) Distributions of viral genome lengths from high (blue) and low (red) contamination regions of the ORR subsurface. (B) Gene-sharing network of ORR viral genomes (red nodes = originating from high contamination regions and blue nodes= originating from low contamination regions) alongside a reduced set, based on clustering with ORR genomes, of ViralRefSeq genomes (grey nodes). Edges represent shared gene content. The shapes of the ORR viral genome node shapes indicate if the genome originated from sediment cores (square), groundwater (circle), or both sample types (triangle). (C) Bar graphs show vCONTACT (left) clustering status and PhaTYP-predicted lifestyles (right) for viral genomes.

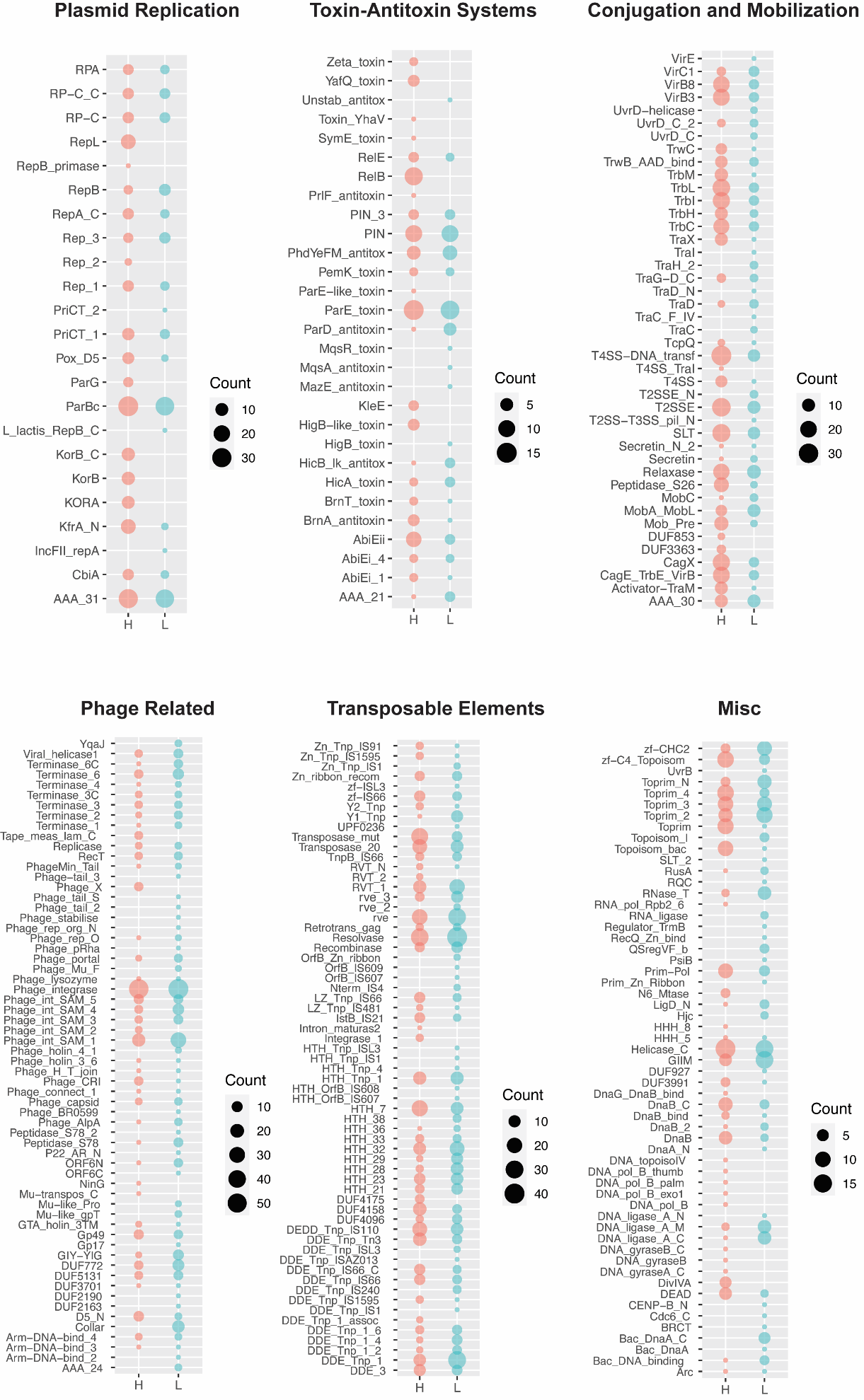

**Figure S2. MGE-related Pfams on unclassifiable circular elements.** Counts of MGE-related Pfam domains on the “unclassified” circular elements from high (H, red) and low (L, blue) [U] sample sets.

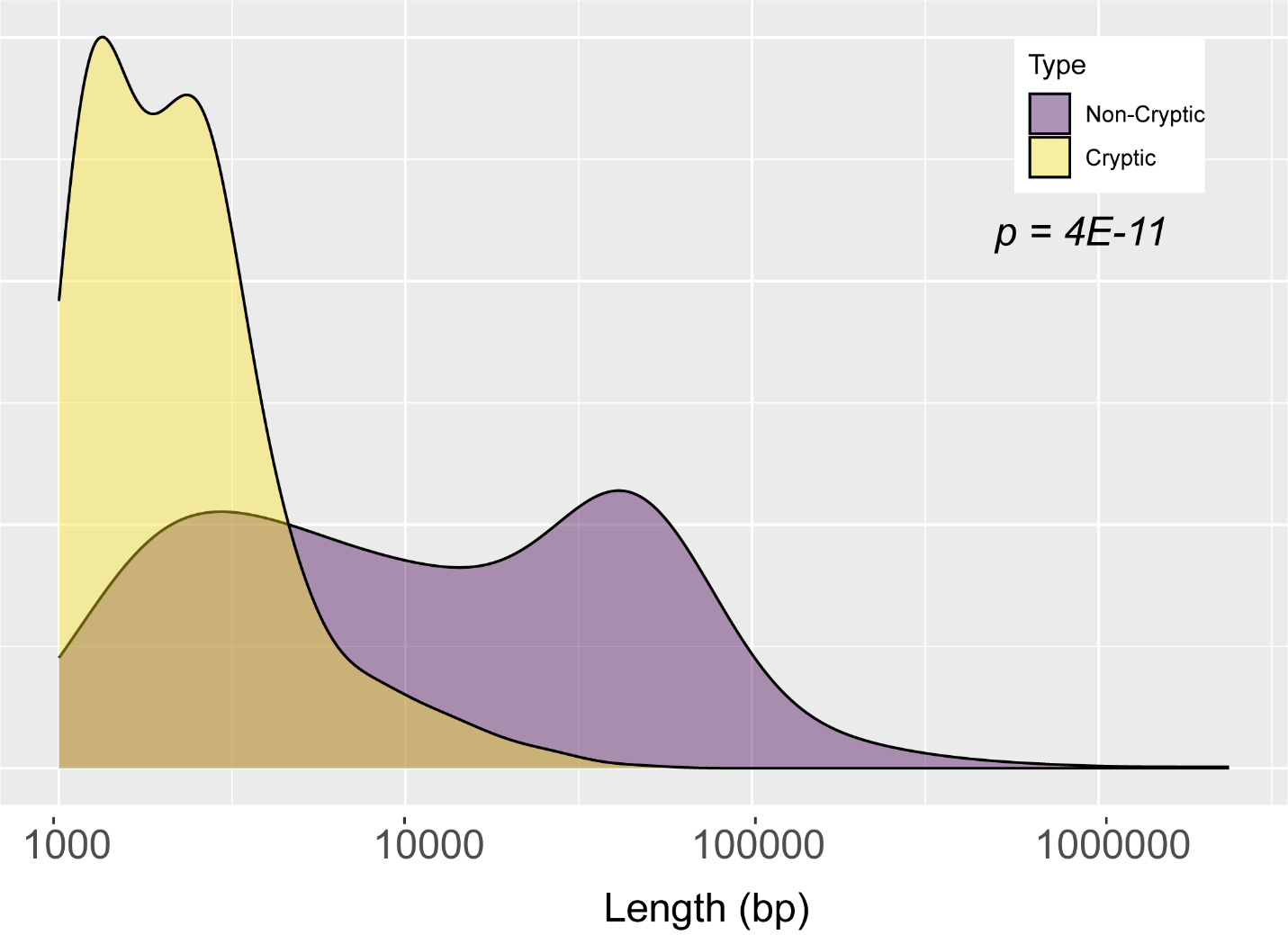

**Figure S3. Length distributions of cryptic vs. non-cryptic MGEs.** Density distributions of the lengths (in bp) of cryptic (yellow) and non-cryptic (purple) MGEs. Data are shown on a Log_10_ x-axis. Statistical comparisons were performed using a two-sided Fisher’s exact test.

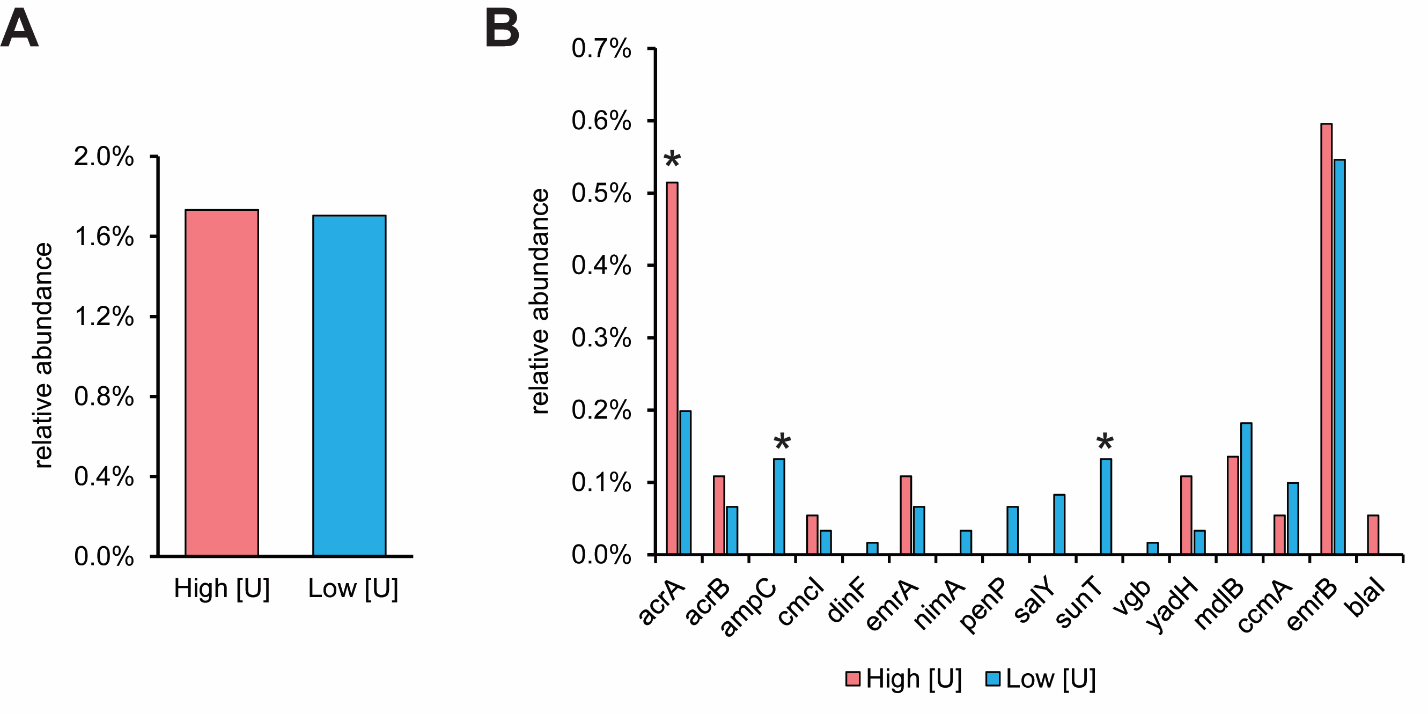

**Figure S4. Analysis of MGE-associated antibiotic resistance genes (ARGs).**

(A) Relative abundance of all identified ARGs normalized against the total annotated CDS in each de-replicated dataset. (B) Relative abundance of individual ARGs normalized against the total annotated CDS in each de-replicated dataset. All statistical comparisons for this figure were performed using a two-sided Fisher’s exact test (* p < 0.05; all others are not significant).

|  | **High [U] Count** | **High [U] Abundance** | **Low [U] Count** | **Low [U] Abundance (%)** | **p-value** |
| --- | --- | --- | --- | --- | --- |
| acrA | 19 | 0.5% | 12 | 0.2% | 0.0157 |
| acrB | 4 | 0.1% | 4 | 0.1% | n.s. |
| ampC | 0 | 0.0% | 7 | 0.1% | 0.0474 |
| ccmA | 2 | 0.1% | 2 | 0.0% | n.s. |
| cmcI | 2 | 0.1% | 1 | 0.0% | n.s. |
| dinF | 0 | 0.0% | 1 | 0.0% | n.s. |
| emrA | 4 | 0.1% | 4 | 0.1% | n.s. |
| emrB | 21 | 0.6% | 33 | 0.6% | n.s. |
| mdlB | 5 | 0.1% | 11 | 0.2% | n.s. |
| nimA | 0 | 0.0% | 2 | 0.0% | n.s. |
| penP | 0 | 0.0% | 4 | 0.1% | n.s. |
| salY | 0 | 0.0% | 3 | 0.1% | n.s. |
| sunT | 0 | 0.0% | 8 | 0.1% | 0.026 |
| blaI | 1 | 0.0% | 0 | 0.0% | n.s. |
| yadH | 4 | 0.1% | 0 | 0.0% | 0.0237 |
| **Total** | **62** | **1.7%** | **92** | **1.7%** | **n.s.** |

**Table S13. Abundance of ARG among non-cryptic circular elements.** Values were re-calculated for sample subset that only included: (1) predicted viruses, (2) circular elements with similarity to known plasmids, and (3) circular elements carrying MGE-related domains. P-values were determined using a two-sided Fisher’s exact test (n.s. is p > 0.05).

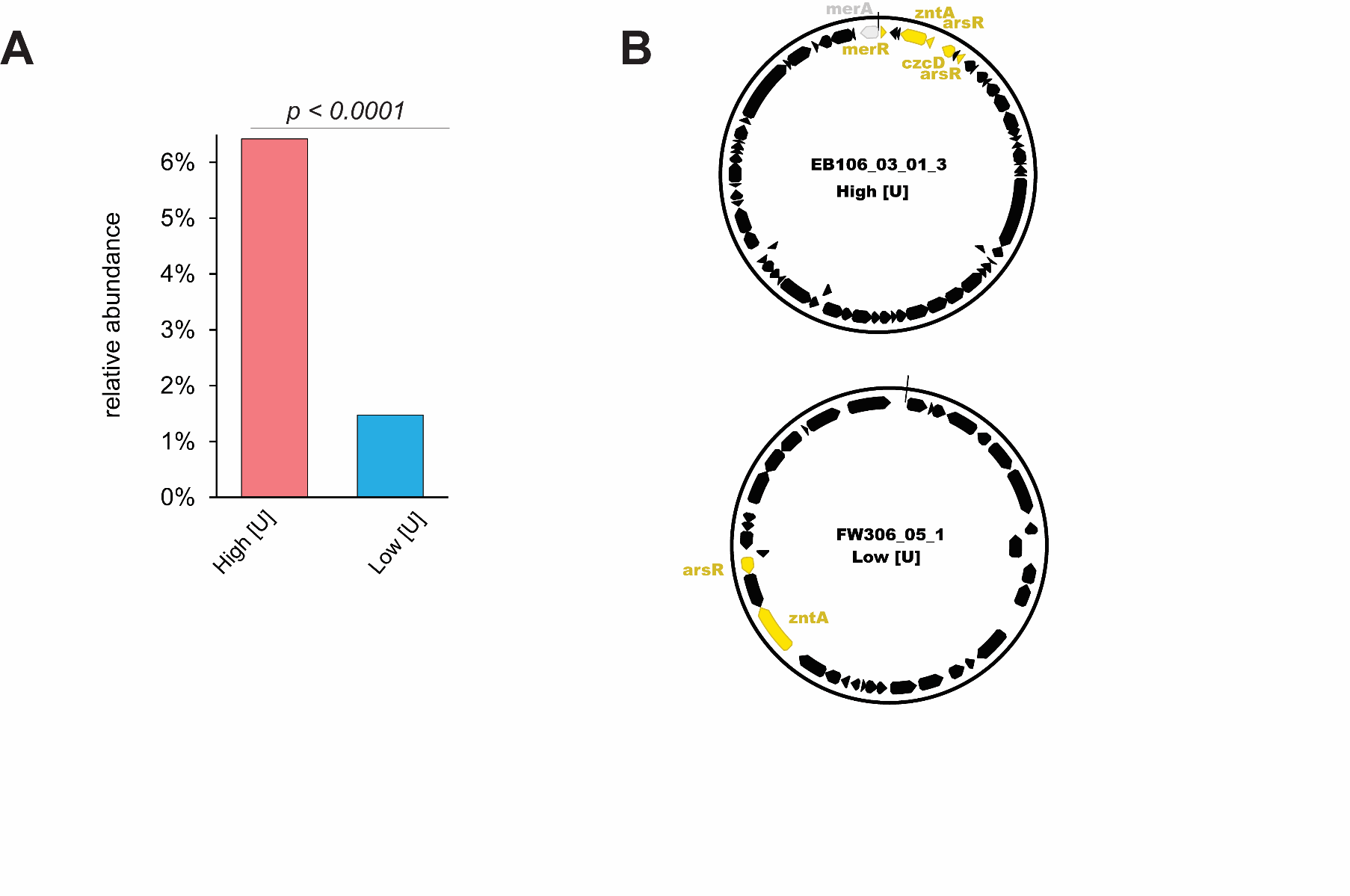

**Figure S5. Analysis of MGE HMRG content.** (A) Relative abundance of all identified HMRGs (see **Table S10** for complete list) normalized against the total annotated CDS in each de-replicated dataset. Statistical comparisons were performed using a two-sided Fisher’s exact test. (B) Maps of two representative plasmids carryign HMRGs from high and low [U] regions of the ORR.

|  | **High [U] Count** | **High [U] Abundance** | **Low [U] Count** | **Low [U] Abundance (%)** | **p-value** |
| --- | --- | --- | --- | --- | --- |
| arsA | 4 | 0.1% | 1 | 0.0% | n.s. |
| acr3 | 0 | 0.0% | 3 | 0.1% | n.s. |
| arsB | 4 | 0.1% | 2 | 0.0% | n.s. |
| arsC | 2 | 0.1% | 4 | 0.1% | n.s. |
| arsR | 14 | 0.4% | 21 | 0.4% | n.s. |
| chrA | 16 | 0.4% | 1 | 0.0% | <0.0001 |
| chrB1 | 12 | 0.3% | 1 | 0.0% | 0.0001 |
| COG2847 | 0 | 0.0% | 1 | 0.0% | n.s. |
| COG3350 | 2 | 0.1% | 0 | 0.0% | n.s. |
| copC | 6 | 0.2% | 1 | 0.0% | 0.017 |
| copZ | 18 | 0.5% | 4 | 0.1% | <0.0001 |
| cusA | 22 | 0.6% | 4 | 0.1% | <0.0001 |
| cusF | 9 | 0.3% | 4 | 0.1% | 0.0425 |
| czcD | 20 | 0.6% | 2 | 0.0% | <0.0001 |
| czcO | 2 | 0.1% | 3 | 0.1% | n.s. |
| fieF | 3 | 0.1% | 2 | 0.0% | n.s. |
| merA | 20 | 0.6% | 4 | 0.1% | <0.0001 |
| merC | 3 | 0.1% | 0 | 0.0% | n.s. |
| merE | 8 | 0.2% | 0 | 0.0% | 0.0006 |
| merR | 34 | 1.0% | 12 | 0.2% | <0.0001 |
| pcoB | 9 | 0.3% | 2 | 0.0% | 0.0092 |
| pcoD | 3 | 0.1% | 1 | 0.0% | n.s. |
| tehB | 0 | 0.0% | 3 | 0.1% | n.s. |
| terC | 0 | 0.0% | 1 | 0.0% | n.s. |
| rcnA | 1 | 0.0% | 0 | 0.0% | n.s. |
| zntA | 17 | 0.5% | 9 | 0.2% | n.s. |
| **Total** | **229** | **6.4%** | **86** | **1.6%** | **<0.0001** |

**Table S14. Abundance of HMRG among non-cryptic circular elements.** Values were re-calculated for sample subset that only included: (1) predicted viruses, (2) circular elements with similarity to known plasmids, and (3) circular elements carrying MGE-related domains. P-values were determined using a two-sided Fisher’s exact test (n.s. is p > 0.05).
